# Harnessing Immunotherapy to Enhance the Systemic Anti-Tumor Effects of Thermosensitive Liposomes

**DOI:** 10.1101/2022.08.29.505721

**Authors:** Maximilian Regenold, Xuehan Wang, Kan Kaneko, Pauric Bannigan, Christine Allen

## Abstract

Chemotherapy plays an important role in debulking tumors in advance of surgery and/or radiotherapy, tackling residual disease, and treating metastatic disease. In recent years many promising advanced drug delivery strategies have emerged that offer more targeted delivery approaches to chemotherapy treatment. For example, thermosensitive liposome mediated drug delivery in combination with localized mild hyperthermia can increase local drug concentrations resulting in a reduction in systemic toxicity and an improvement in local disease control. However, the majority of solid tumor associated deaths are due to metastatic spread. A therapeutic approach focused on a localized target area harbors the risk of overlooking and undertreating potential metastatic spread. Previous studies reported systemic, albeit limited, anti-tumor effects following treatment with thermosensitive liposomal chemotherapy and localized mild hyperthermia. This work explores the systemic treatment capabilities of a thermosensitive liposome formulation of the vinca alkaloid vinorelbine in combination with mild hyperthermia in an immunocompetent murine model of rhabdomyosarcoma. This treatment approach was found to be highly effective at heated, primary tumor sites. However, it demonstrated limited anti-tumor effects in secondary, distant tumors. As a result, the addition of immune checkpoint inhibition therapy was pursued to further enhance the systemic anti-tumor effect of this treatment approach. Once combined with immune checkpoint inhibition therapy, a significant improvement in systemic treatment capability was achieved. We believe this is one of the first studies to demonstrate that a triple combination of thermosensitive liposomes, localized mild hyperthermia, and immune checkpoint inhibition therapy can enhance the systemic treatment capabilities of thermosensitive liposomes.

**Graphical abstract:** 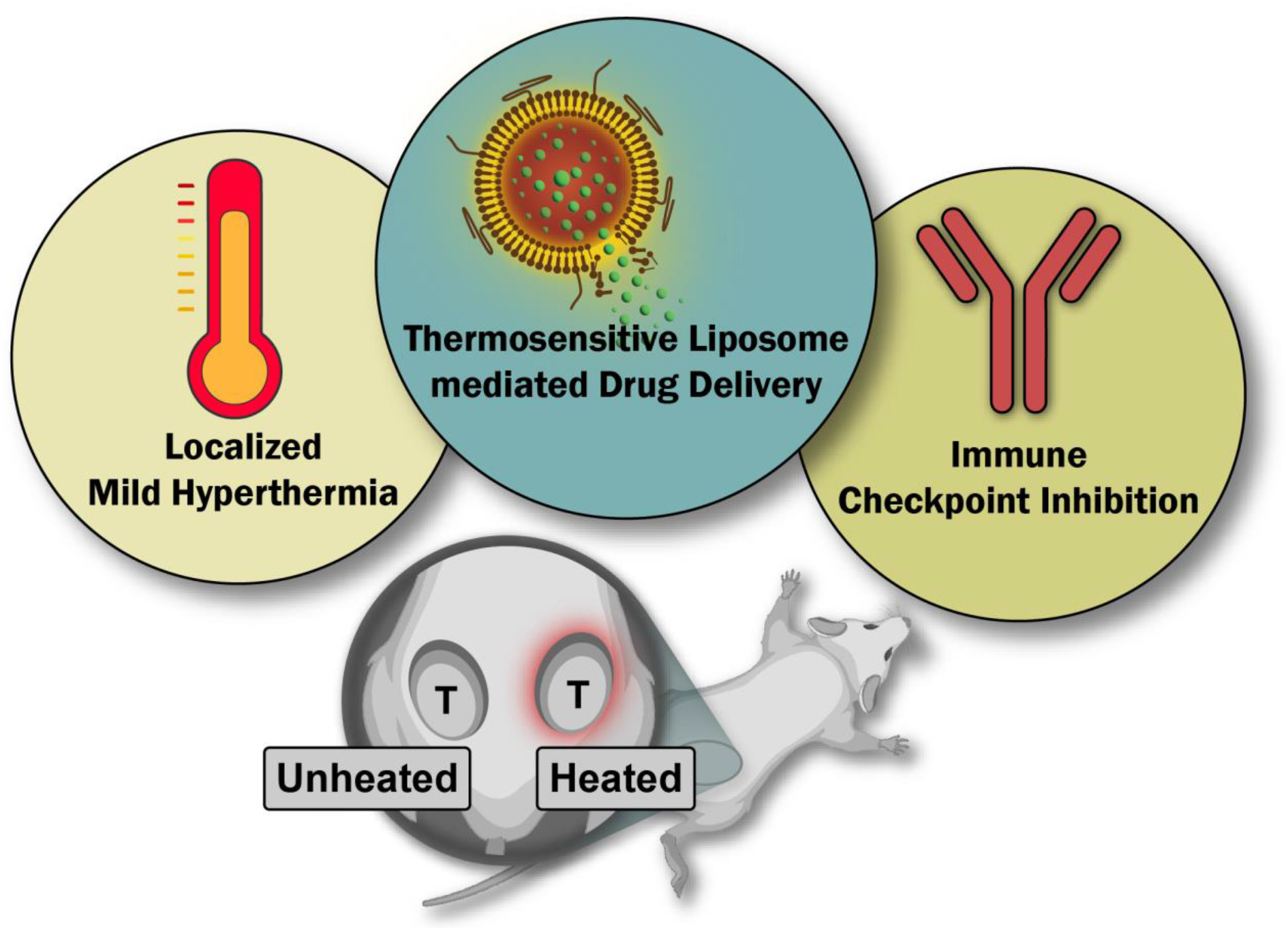

## 1. Introduction

Surgery and radiotherapy remain the cornerstones of local cancer therapy [1]. While chemotherapy is commonly used as a systemic treatment approach to debulk tumors in preparation for surgery and/or radiotherapy, treat residual disease and reach metastatic disease sites. Incomplete local disease control can lead to recurrence at the primary site [2]. Such instances of local recurrence are known to occur in a number of different cancers, including non-small cell lung cancer, colorectal cancer, malignant pleural mesothelioma and rhabdomyosarcoma (RMS) [2]. In the case of RMS, up to one third of patients experience recurrence and 70-80 % of these cases occur at the primary disease site [3]. Reducing the likelihood of disease recurrence requires more aggressive therapeutic strategies. However, these are often accompanied by more severe adverse effects. Consideration of potential long-term adverse effects can play an important role treatment selection, in particular in cancers such as RMS, where the majority of cases present in patients under 10 years of age [4,5].

One option to improve local disease control is to employ drug delivery strategies, such as nanoparticles, that provide triggered drug release and result in an increase in tumor drug concentrations. These systems are administered systemically and designed to release their content in the presence of a specific trigger. To this end, delivery platforms exploiting both internal (e.g., pH or hypoxia) and/or external (e.g., temperature or ultrasound) triggers to stimulate drug release have been developed [6]. This includes thermosensitive liposomes, initially developed by Yatvin et al. over four decades ago with the goal of improving upon passively targeted liposomes [7]. Since then, various chemotherapy drugs have been formulated as thermosensitive liposomes with doxorubicin being the most frequently explored compound. In many cases, these triggered delivery systems have been evaluated in combination with localized heating and have afforded promising improvements in treatment efficacy in animal models of local disease [8–10]. ThermoDox®, a thermosensitive liposome formulation encapsulating doxorubicin, is the only formulation that has been evaluated in clinical trials [11] However, as discussed elsewhere, ThermoDox has faced significant setbacks in its clinical development, in particular when evaluated in combination with radiofrequency ablation (temperatures > 50 °C) [11]. Nonetheless, several pre-clinical and clinical studies have demonstrated the potential of this treatment approach in combination with mild hyperthermia (temperatures of 39-45 °C), with a number of studies achieving significant improvements in the drug’s therapeutic index [12–14].

Currently the majority of cancer associated deaths are due to metastatic spread to secondary sites [15]. An important consideration regarding the treatment of metastatic disease is that achieving local control (e.g., via surgery or radiotherapy) is recognized as a pivotal step in reducing the probability of metastatic spread in the first place [16]. However, due to challenges in detecting metastatic sites, many patients are estimated to harbor undetected metastases at the time of initial surgery [16]. Thus, despite the need for local control in cancer therapy, there is clearly a need for a treatment approach that offers both local as well as systemic anti-tumor treatment capabilities. This raises the question: can systemically administered thermosensitive liposomes, that are designed to improve local disease control, also treat metastatic disease?

The concept of a local treatment approach inducing a systemic anti-tumor effect is commonly referred to as the ‘abscopal effect’. The first mention of this effect dates back to a 1953 publication on whole body irradiation, where Robin Mole asked the question “has irradiation of a mammal an effect at a distance from the volume irradiated?” [17]. However, to this day, the potential systemic effects of ionizing radiation remain highly questionable with few documented occurrences over the past several decades [18]. In fact, only 46 clinical cases of radiotherapy-induced abscopal effects were reported between 1969 and 2014 [19]. Yet, the emergence of immune checkpoint inhibition (ICI) therapy has led to a significant increase in pre-clinical and clinical reports of the abscopal effect [20]. While the exact mechanism remains to be fully understood, it has been shown that the immune system plays a key role [21]. Radiation induced cell damage leads to the production and release of antigens, damage-associated molecular patterns (DAMPs), as well as cytokines, and as a consequence stimulates a systemic cytotoxic T cell based anti-tumor immune effect [22]. However, the scarcity of the abscopal effect indicates that this immune response may be challenged by immunosuppressive signals, including T cell inhibitory receptors such as CTLA-4 and PD-1. Indeed, there is evidence that the addition of ICI therapy (e.g., anti-CTLA-4 or anti-PD-1 antibodies) can increase tumor-targeted cytotoxic T cell activity and therefore boost the abscopal effect of radiotherapy [20]. This approach has since shown promising results in several clinical trials [23].

More recently, the abscopal effect has been referenced in connection with other localized treatment strategies that induce a systemic effect (e.g., intratumoral chemotherapy, cryotherapy, ablation, and mild hyperthermia) [24]. If the therapeutic intervention is confined to a specific treatment volume, the resulting systemic effects can generally be described as abscopal. Localized mild hyperthermia treatment (39-43 °C) is known to stimulate the immune system and enhance anti-tumor immune effects [25,26]. As far back as 2009, Skitzki et al. highlighted the potential of mild hyperthermia as a non-invasive, non-toxic, and readily available treatment modality that could be used as an adjuvant with existing immunotherapy approaches [26]. In the clinic, localized mild hyperthermia is commonly applied in combination with radiotherapy focusing on mild hyperthermia as an approach to boost radiation induced abscopal effects [27,28]. More recently, the immune stimulating effects of mild hyperthermia treatment are under investigation in combination with immunotherapy (NCT03757858, NCT03393858).

In the last ten years, chemotherapy has increasingly been recognized for its potential impact on the immune system [29,30]. The combination of chemotherapy with ICI has been suggested as a promising approach to produce systemic anti-cancer immunity, similar to the aforementioned concept of ICI boosting radiation induced abscopal effects [31]. Thermosensitive liposome mediated chemotherapy could be of particular interest since it not only targets drug specifically to the tumor but also comes with the immune stimulating effects of localized mild hyperthermia treatment. Importantly, despite the heating being applied locally, this delivery approach does not preclude systemic drug exposure. Thus, off-target anti-tumor effects should not be referred to as abscopal effects but rather as anenestic tumor effects [32]. Here, enestic describes a tumor lesion exposed to intratumoral treatment (here externally triggered intravascular release of chemotherapeutics) and in consequence anenestic refers to ‘non-injected’ lesions.

The study presented herein is one of the first to combine ICI (i.e., PD-1 blockade alone) with mild hyperthermia-triggered thermosensitive liposome mediated chemotherapy. First, we explored the treatment efficacy of our previously developed thermosensitive liposome formulation encapsulating vinorelbine (ThermoVRL) in an immunocompetent model of RMS. We then assessed potential anenestic tumor effects of this treatment approach in a bilateral tumor model. And lastly, determined if the addition of ICI in the form of PD-1 blockade therapy enhances systemic anti-tumor effects. Overall, the results demonstrate the significant local treatment effect of ThermoVRL, as well as a limited effect on distant tumors, when combined with localized mild hyperthermia. Interestingly, the addition of PD-1 blockade therapy was found to significantly boost anenestic tumor effects. Moreover, we were able to demonstrate abscopal effects of localized mild hyperthermia alone with the addition of PD-1 blockade therapy. In summary, this study shows that the combination of ICI and mild hyperthermia-triggered thermosensitive liposome mediated chemotherapy has tremendous potential as a multimodal treatment approach for the management of local and metastatic disease.

## 2. Materials and Methods

### 2.1. Materials

Vinorelbine tartrate (VRL) was obtained from Selleck Chemicals (Houston, TX, USA). Anti-PD-1 antibody was purchased from Bio X Cell (RMP1-14, Bio X Cell, Lebanon, NH, USA). Sodium sucrose octasulfate (Na_8_SOS) was purchased from Toronto Research Chemicals (North York, ON, Canada). 1,2-Dipalmitoyl-sn-glycero-3-phosphocholine (DPPC), N-(carbonyl-methoxypolyethyleneglycol 2000)-1,2-distearoyl-sn-glycero-3-phosphoethanolamine (PEG_2k_-DSPE), 1-stearoyl-2-lyso-sn-glycero-3-phosphocholine (lyso-SPC, MSPC) were obtained from Corden Pharma (Plankstadt, Germany). Triethylamine (TEA), Dowex® 50WX8-200, fetal bovine serum (FBS), penicillin and streptomycin (P/S), and 2-mercaptoethanol were purchased from Sigma-Aldrich (Oakville, ON, Canada). M3–9-M cells were kindly provided by Dr. Crystal Mackall (Stanford University). Cell culture medium was purchased from Life Technologies (Burlington, ON, Canada).

### 2.2. Cell culture and cytotoxicity evaluation

The murine RMS cell line M3–9-M was grown in RPMI 1640 supplemented with 10 % FBS, 1 % P/S, 1 % L-Glutamine, 1 % NEAA, 1 % sodium pyruvate and 50 µM 2-mercaptoethanol. Cells were kept at 37 °C with 5 % CO_2_, unless otherwise indicated. Cells were seeded in 96-well plates at 100 cells per well followed by treatment with VRL after 24 h of incubation. Cell viability was then assessed after 72 h, using an acid phosphatase assay. Specifically, 2 mg/mL phosphatase substrate p-nitrophenylphophate was added to cells for 1 h followed by the addition of 0.1 N NaOH and UV absorbance measurement at 405 nm. Half maximal inhibitory concentration (IC_50_) values were obtained using GraphPad Prism 6.0 (GraphPad Software, San Diego, CA, USA) by fitting the data with a 4-parameter sigmoidal dose response curve. To evaluate the effect of mild hyperthermia on VRL cytotoxicity, cells were incubated at 42 °C for one hour followed by 71 h at 37 °C.

### 2.3. Liposome preparation

Thermosensitive liposomes loaded with VRL were prepared as previously described [33]. In brief, a thin lipid film was prepared by dissolving DPPC, MSPC, and PEG_2k_-DSPE at a molar ratio of 86/10/4 in chloroform. This solution was then dried using a rotary evaporator. The film was then hydrated with a TEA_8_SOS solution (0.22 M sulfate group concentration) to a total lipid concentration of 125 mM. The liposomes were then extruded (Lipex Extruder, Northern Lipids, Vancouver, BC, Canada) three times through double stacked track-etch polycarbonate membranes with a 200 nm pore size (Whatman Inc., Clifton, NJ, USA). Following this, the resulting liposomes were extruded a further 10 times through 100 nm pore size membranes. Extruded liposomes were then chilled on ice for 10 min prior to over-night dialysis in HBS (20 mM HEPES, 150 mM sodium chloride, pH 6.5). VRL was loaded at 30 g VRL/mol lipid by incubating the liposomes with drug at 35 °C for 60 min. Liposomes were subsequently chilled on ice for 10 min, purified by overnight dialysis in HBS and concentrated via tangential flow filtration using a polysulfone MicroKros® filter (Spectrum, Rancho Dominguez, CA, USA).

### 2.4. Tumor model and localized mild hyperthermia treatment

All animal studies were conducted in accordance with the guidelines of the Animal Care Committee of the University Health Network (UHN, Toronto, ON, Canada). An aliquot of 2 × 10^6^ M3–9-M cells was injected subcutaneously into either the right flank only (**Schematic 1**A: single tumor study), or into the right and left flanks (**Schematic 1**B: double tumor study) of 6-8-week-old male C57BL/6 mice to develop ectopic syngeneic tumors. Tumor volumes were calculated as 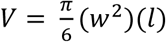 from measurements of tumor width (*w*) and length (*l*). Treatments were started once one tumor reached a volume > 90 mm³. In the bilateral tumor model, the larger of the two tumors was considered the primary tumor and thus determined to be the site of mild hyperthermia (i.e., HT). This tumor was pre-heated to 42.5 °C for 5 min. Following this 5 min period, intravenous tail vein injection of ThermoVRL or saline was administered under constant heating, and heating was continued for another 20 min post treatment. The contralateral tumor remained unheated. A laser-based heating system developed by Dou et al., in combination with a centrally placed single-point temperature probe, was used to achieve the desired tumor temperatures [34]. 200 µg anti-PD-1 antibody was administered intraperitoneally (i.p.) twice per week for 35 days. This treatment was started one day prior to ThermoVRL and/ or mild HT administration. Tumor dimensions of > 15 mm in any dimension, or a body weight loss > 20 %, were selected as ethical endpoints. The bilateral tumor study (**Schematic 1**B) was concluded as soon as no palpable tumors were detectable since previous studies in our laboratories demonstrated a very durable treatment response with no tumor re-growth past a specific study timepoint.

### 2.5 Statistical analysis

Statistical analysis was performed using SSPS Statistics 28.0 (IBM, Armonk, NY, USA). Animal weights on specific treatment days were compared by one-way ANOVA with Bonferroni *post hoc* testing. Survival differences were calculated and compared with a log-rank test and a Bonferroni correction for multiple comparisons. Tumor volumes of different treatment groups were compared by unpaired *t*-test.

## 3. Results

ThermoVRL was prepared as previously described [33]. Prior to *in vivo* administration, the drug loaded liposomes were characterized in terms of size, zeta potential, melting phase transition temperature, as well as drug loading and the data were in agreement previously published results.

The studies were conducted in an immunocompetent murine model of RMS [35]. Specifically, M3–9-M cells were used to grow subcutaneous tumors in male C57BL/6 mice. As shown in **Schematic 1**A, tumors were heated to mild hyperthermia temperatures (HT; 42.5 ± 0.5°C) for 5 min prior to intravenous administration of free VRL or ThermoVRL [36]. Pre-heating was performed to ensure maximum intravascular drug release prior to liposome clearance from the systemic circulation. Localized heating was then continued for another 20 min.

**Schematic 1:**
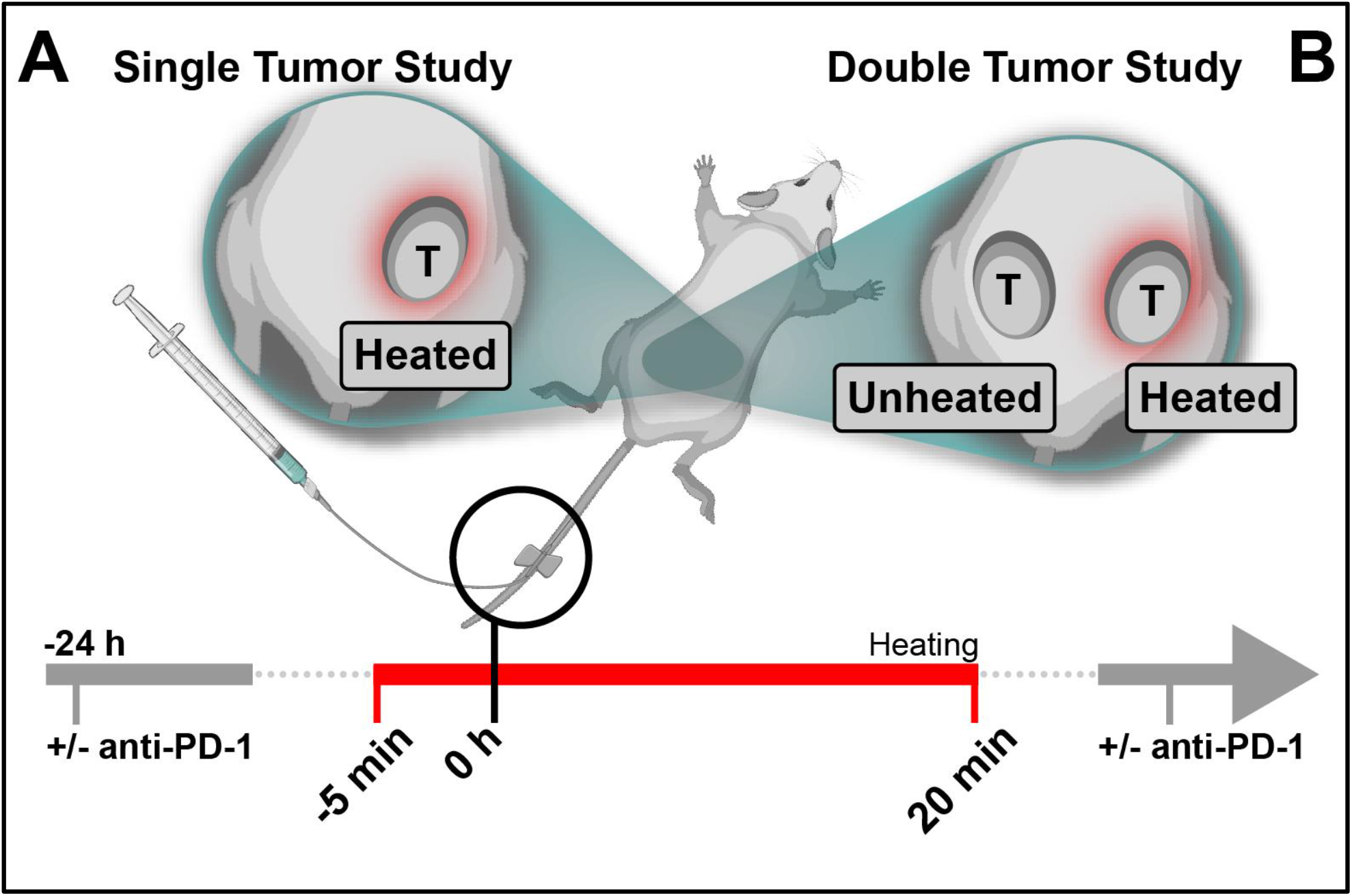
Overview of the design of both the single tumor and double tumor studies. (A) To determine the treatment efficacy of VRL encapsulated in thermosensitive liposomes (ThermoVRL), single M3–9-M tumor bearing male C57BL/6 mice were treated with free VRL or ThermoVRL in combination with mild hyperthermia (42.5 °C). Tumors were pre-heated for 5 min prior to intravenous treatment administration followed by another 20 min of heating. (B) Distant effects of this treatment approach were determined using a bilateral tumor model where one tumor was subjected to the same heating protocol as described in (A), while the second tumor remained unheated. As part of study (B), the addition of immune checkpoint inhibition therapy (+ anti-PD-1) was pursued to enhance distant anti-tumor effects. Specifically, animals treated with ThermoVRL and localized mild hyperthermia received anti-PD-1 treatment one day prior to chemotherapy administration. Bi-weekly anti-PD-1 treatment was then continued for a total of five weeks.

To evaluate the efficacy of ThermoVRL (+/- HT) in this immunocompetent animal model, controls of saline (+/- HT) and free VRL (+/- HT) were used. Hyperthermia is known to have direct cytotoxic effects particularly on cancer cells [37]. However, this cytotoxicity depends on both the temperature and the duration of heating [38]. In the studies presented, localized HT treatment (compared to saline alone) did not increase median survival (11 ± 1 days versus 10 ± 1 days, respectively). The heating protocol employed was designed to effectively trigger intravascular drug release from thermosensitive liposomes. Higher temperatures or extended heating times would likely be required to result in hyperthermia mediated cytotoxicity. Treatment with free VRL at 10 mg VRL/kg body weight increased the median survival time (20 ± 10 days) in comparison to the administration of saline (10 ± 1 days), albeit this increase was not statistically significant. However, the addition of HT further improved survival times of animals treated with free VRL (26 ± 5 days), leading to a significant difference in comparison to saline treated animals (p = .04). In fact, 2 out of the 5 mice treated with free VRL + HT went into complete remission with no regrowth of tumors prior to the end of the study (120 days). Interestingly, in our previous study which explored the therapeutic effect of VRL and liposomal formulations of VRL in an immunocompromised model of RMS, we did not observe a benefit associated with the addition of HT to free drug treatment [39].

As expected, adding HT to ThermoVRL treatment significantly improved median survival times relative to treatment with ThermoVRL alone (14 ± 1 days versus > 120 days; p < .03). In fact, all animals treated with ThermoVRL in combination with HT went into complete remission and no palpable tumors were detected for the remainder of the study.

The body weight of mice was recorded as a measure of treatment toxicity. A decrease in body weight was observed in animals administered VRL (free or ThermoVRL) immediately following treatment, but animals appeared to recover quickly with weights reaching baseline levels within 10 days post treatment.

Treatment of localized tumors with ThermoVRL + HT in this immunocompetent model of RMS proved to be highly effective. In order to investigate the systemic treatment capabilities of this approach, a bilateral tumor model was employed (**Schematic 1**B). M3-9-M cells were injected subcutaneously into both the left and right hind limbs of each mouse. Following tumor development, the tumor with the larger volume was considered the ‘primary’ tumor, while contralateral tumors were used as a proxy for metastatic disease. HT was applied locally to the primary tumor only, and the contralateral tumor remained unheated. The average volume of primary (i.e., heated) tumors was 162 ± 72 mm³ compared to 85 ± 48 mm³ for the contralateral (i.e., unheated) tumors (p < .001). Control groups for this study included animals bearing bilateral tumors that received (i) no treatment (untreated); (ii) intraperitoneal (i.p.) PD-1 blockade therapy (anti-PD-1); (iii) intravenous saline and HT (saline + HT).

Interestingly, as shown in **Figure 2**, the saline + HT group exhibited significantly smaller primary (i.e., heated) tumor volumes at day 11 compared to tumors of animals in the untreated group (p = .03). However, no difference in tumor volumes was observed between the contralateral (i.e., unheated) tumors of animals in the saline + HT group and tumors of animals in the untreated group. This resulted in no difference in median survival times between these treatment groups (**Figure 4**). This indicates that tumor growth inhibition due to HT was limited to the primary tumor and did not yield an abscopal effect. As expected, treatment with ThermoVRL + HT led to complete primary tumor remission in 9 out of 10 animals (**Figure 2**). Volumes of contralateral tumors were significantly reduced compared to corresponding tumors of the saline + HT control group (on day 11, p < .001). The limited growth inhibition of contralateral (i.e., unheated) tumors in animals treated with ThermoVRL + HT led to animals reaching ethical endpoints within 37 days. Nonetheless, median survival times of animals treated with ThermoVRL + HT were significantly prolonged compared to animals treated with saline +HT or untreated animals (p < .01). In fact, the median survival time was found to be similar to that of mice receiving free drug treatment in the single tumor study (**Schematic 1**A) (25 ± 1 days and 20 ±10 days, respectively). Intravascular drug release triggered at the target site is known to increase the amount of drug delivered to the tumor [34,40], however, a significant amount of released drug can access the systemic circulation [10]. It appears that drug molecules that enter the systemic circulation (following ThermoVRL + HT treatment) do significantly influence the growth of contralateral tumors (**Figure 2**). However, the anenestic tumor effects of ThermoVRL + HT stand in stark contrast to the effect observed on primary, heated tumors (i.e., where complete remission was observed). This does highlight the tremendous potential of ThermoVRL + HT to provide local anti-tumor effects, but it also demonstrates the limited effects of this treatment approach on secondary, unheated tumors.

**Figure 1:**
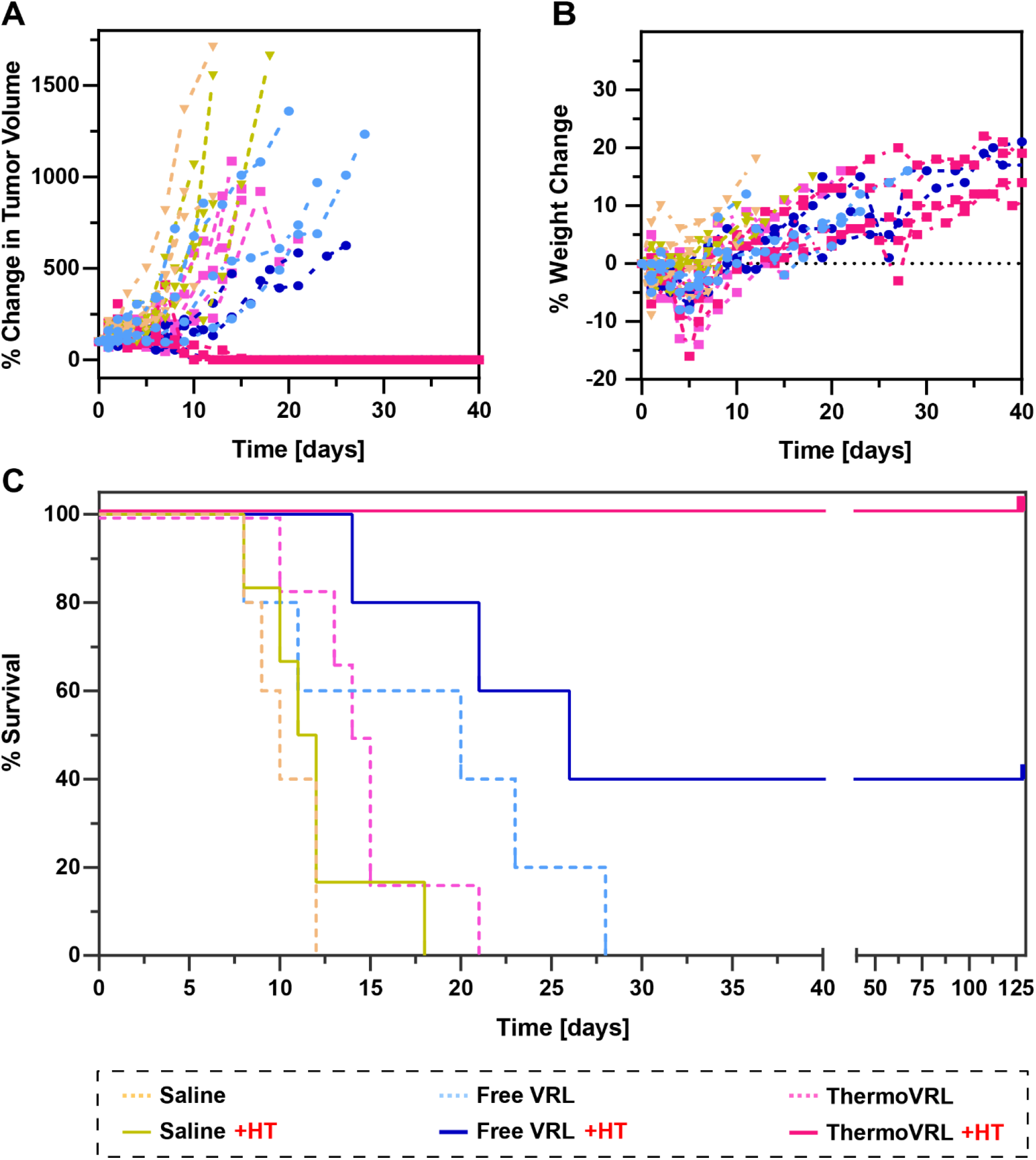
*In vivo* efficacy study comparing the different treatment groups with and without the addition of mild hyperthermia (HT) localized to the tumor site. Male C57BL/6 mice bearing subcutaneous M3–9-M tumors were treated once on day 0 with 10 mg VRL/kg body weight. A) Tumor volume change, b) body weight change, and c) Kaplan-Meier survival analysis are shown here. The addition of localized HT to treatment with thermosensitive liposomes loaded with VRL (ThermoVRL) improved median survival times nearly 9-fold (p < .03). n ≥ 5 mice per group.

**Figure 2:**
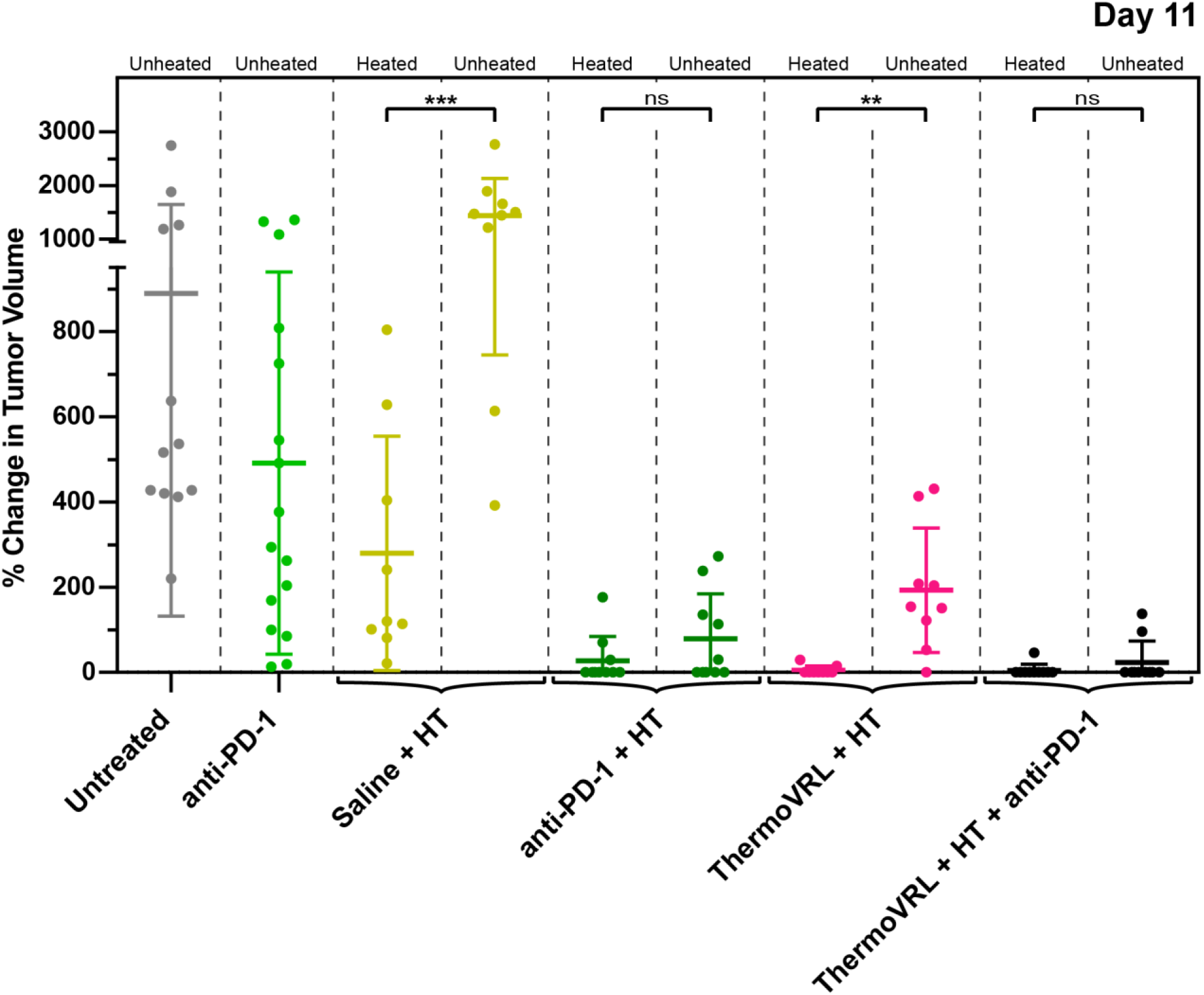
Volume change of bilateral M3–9-M tumors in male C57BL/6 mice on day 11 post treatment. Thermosensitive liposomal vinorelbine (ThermoVRL) was administered at 10 mg/ kg body weight. Saline and ThermoVRL were administered once intravenously on day 0, while immune checkpoint inhibition (ICI) therapy via anti-PD-1 antibodies was administered intraperitoneally on day -1 and continued twice per week for a total of 5 weeks. The primary tumor was subjected to mild hyperthermia (HT; 42.5 °C, 25 min) treatment (i.e., heated), while the contralateral tumor remained unheated (i.e., unheated). Addition of ICI therapy to mild HT treatment alone resulted in a significant abscopal effect. More importantly, the addition of ICI to ThermoVRL + HT provided significant anenestic tumor effects resulting in complete contralateral tumor remission in 8 out of 10 mice on day 11 post treatment. n ≥ 8 mice per group.

**Figure 3:**
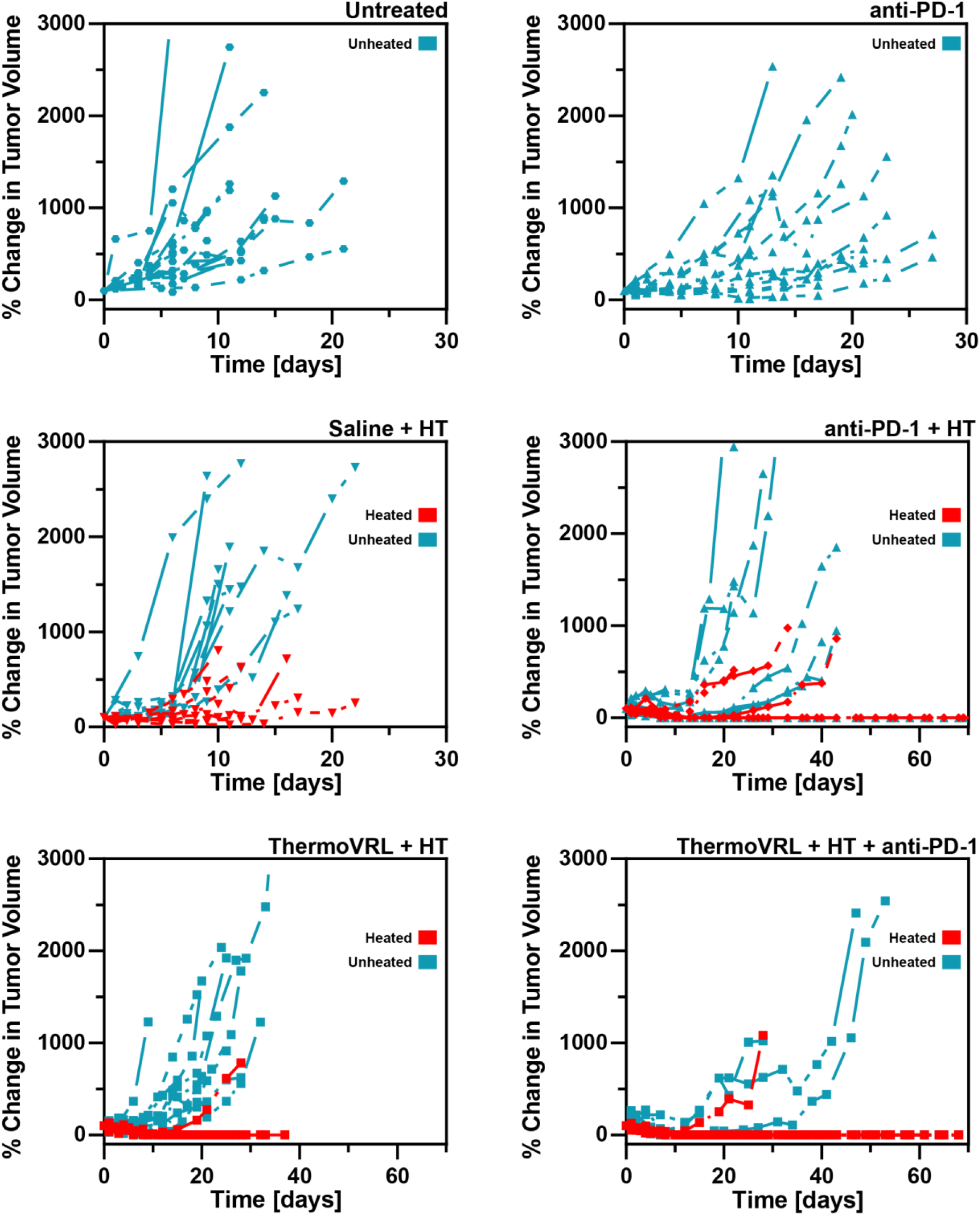
Male C57BL/6 mice bearing bilateral, subcutaneous, M3–9-M tumors received either no treatment (untreated), intraperitoneal PD-1 blockade therapy (anti-PD-1), intravenous saline (saline), or intravenous thermosensitive liposomal vinorelbine (ThermoVRL) at 10 mg VRL/kg body weight. As indicated (+ HT), several of these treatments were administered simultaneously with mild hyperthermia (42.5 °C, 25 min) localized to the primary tumor. Tumor growth of contralateral (i.e., unheated) tumors is indicated in blue, while growth of primary (i.e., heated) tumors is shown in red. Comparing the tumor growth of animals that received saline + HT and animals treated with anti-PD-1 + HT reveals abscopal effects in the form of slower tumor growth in the contralateral tumors. Only one primary tumor in the ThermoVRL + HT + anti-PD-1 group did not respond to therapy. This triple combination also afforded significantly slower tumor growth of contralateral tumors. n ≥ 8 mice per group.

**Figure 4:**
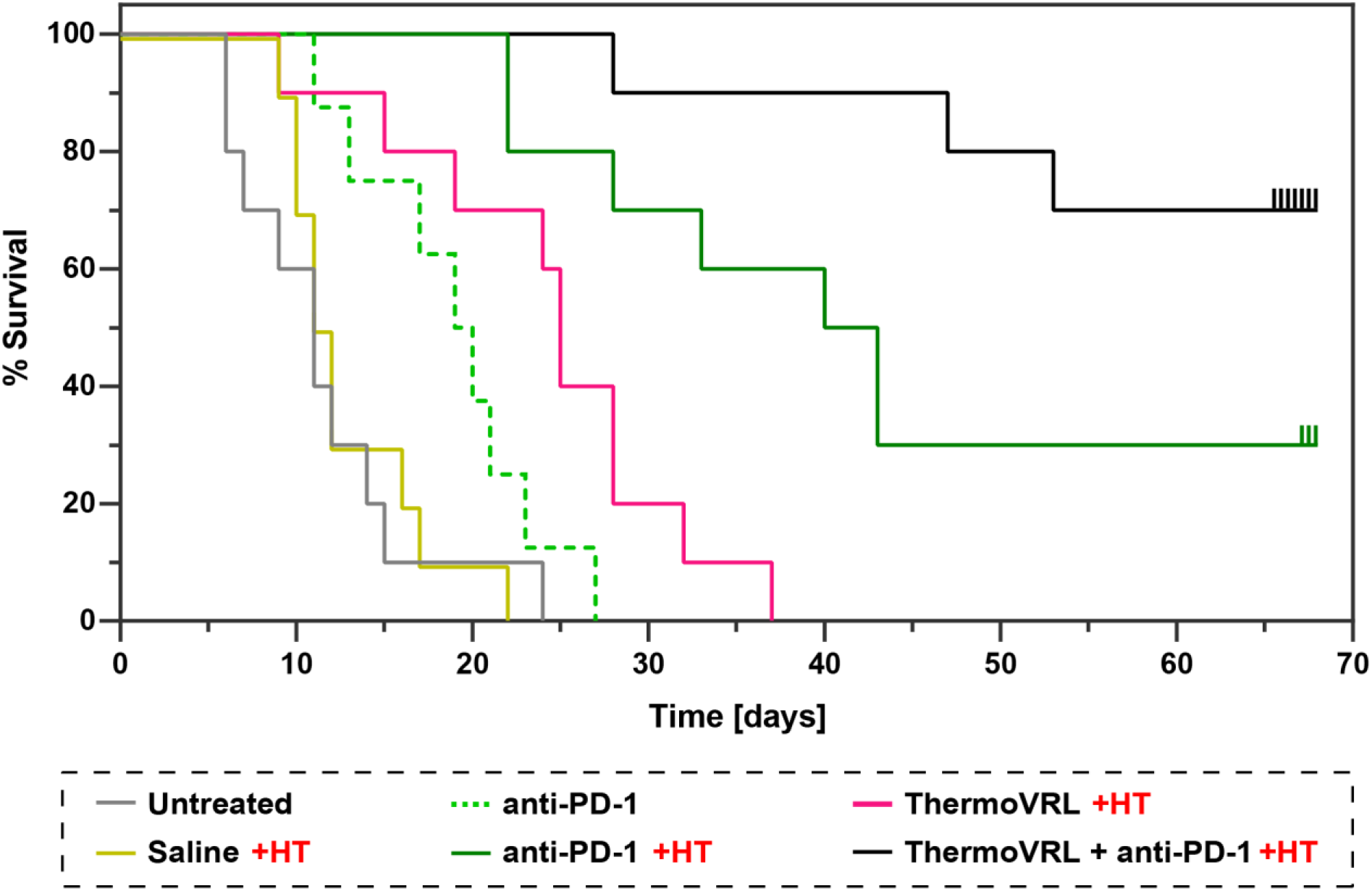
Efficacy study evaluating systemic treatment effects of thermosensitive liposome mediated vinorelbine chemotherapy in combination with localized mild hyperthermia and immune checkpoint inhibition (ICI) therapy. Male C57BL/6 mice bearing bilateral, subcutaneous, M3–9-M tumors received either saline (saline) or thermosensitive liposomes loaded with vinorelbine (ThermoVRL) at 10 mg/kg body weight via intravenous teil vein injection. Primary tumors were heated to mild hyperthermic temperatures (HT) five minutes prior to intravenous treatment administration followed by heating for another 20 min. Contralateral tumors remained unheated. Intraperitoneal administration of 200 µg anti-PD-1 antibody was commenced 24 h prior to treatment with saline, ThermoVRL, or mild HT alone. PD-1 blockade therapy was subsequently administered twice per week for a total of five weeks. The addition of ICI therapy to mild HT (i.e., saline + HT compared to anti PD-1 + HT) increased survival times 3.6-fold (p < .001). The addition of ICI to ThermoVRL (i.e., ThermoVRL + HT compared to ThermoVRL + HT + anti-PD-1) therapy increased median survival times 2.7-fold (p < .001). n ≥ 8 mice per group.

In order to extend beyond the localized treatment constraints of ThermoVRL + HT, a multimodal strategy including immunotherapy was evaluated. Specifically, animals received 200 µg of anti-PD-1 antibody administered i.p. approximately 24 h prior to treatment with ThermoVRL + HT. Anti-PD-1 injections were subsequently continued twice per week for a total of five weeks or until animals reached endpoint. No significant difference in tumor volume between primary (i.e., heated) and contralateral (i.e., unheated) tumors was observed within the anti-PD-1 + HT group on day 11 (**Figure 2**). The tumor volumes for both the primary and contralateral tumors of the anti-PD-1 + HT animals were significantly reduced compared to the anti-PD-1 control (on day 11; p < .01). Thus, indicating that these abscopal effects are stimulated by anti-PD-1 + HT and not anti-PD-1 treatment alone. This combination (i.e., anti-PD-1 + HT) did induce complete remission of both the primary and contralateral tumors in 3 out of 10 animals (compared to 0 out of 8 mice treated with anti-PD-1 only) (**Figure 3**). Moreover, 7 out of 10 of the primary (i.e., heated) tumors in mice receiving anti-PD-1 + HT were undetectable at the time of sacrifice or study termination. Overall, this led to a 2-fold increase in survival time for animals receiving anti-PD-1 + HT compared to anti-PD-1 alone (p < .01).

As shown in **Figure 2**, combining ThermoVRL + HT with PD-1 blockade therapy significantly reduced growth of contralateral (i.e., unheated) tumors compared to treatment with ThermoVRL + HT alone. This led to more than 2.5-fold increase in median survival time (> 68 days versus 25 ± 1 days; p < .001) (**Figure** 4). In fact, for 7 out of the 10 mice, both the primary (i.e., heated) and contralateral (i.e., unheated) tumors went into complete remission with no palpable tumor regrowth for the duration of the study. The increase in median survival time was significantly greater than that resulting from treatment with anti PD-1 alone (19 ± 2 days; p < .001). While the triple combination, ThermoVRL + HT + anti-PD-1, prolonged survival compared to anti-PD-1 + HT almost 1.6-fold, this difference was not found to be statistically significant (> 68 days versus 40 ± 5 days; p = .633).

## 4. Discussion

### Treatment with thermosensitive liposomes and localized mild hyperthermia

Systemic administration of thermosensitive liposomes in combination with localized mild hyperthermia has been proven to be highly effective in the treatment of local disease [41–43]. These improvements are largely attributed to increased accumulation and distribution of drug molecules at the tumor site [44–46]. However, this delivery approach requires heating of the target region as a trigger for drug release. Advances in heating techniques, thermometry, and treatment planning have significantly expanded the types, volumes, and locations of tumors that are amenable to mild hyperthermic heating [47–49]. Thus, expanding the clinical scenarios in which thermosensitive liposomes can be explored. However, any therapeutic approach focused on a localized target risks undertreating metastatic disease. Indeed, metastatic spread continues to be the primary cause of death from cancer [50,51]. This raises the question of how to increase the systemic anti-tumor effects of this treatment approach. Previously, Viglianti et al. demonstrated that a formulation equivalent to ThermoDox was able to reduce tumor growth in secondary, unheated tumors in a bilateral immunodeficient mouse model [10]. The authors concluded that this effect “is most likely due to recirculation of intravascularly released drug” [10]. However, there are several factors that need to be considered prior to generalizing these findings. First, the drug’s pharmacokinetic properties will govern its circulation behaviour and thus impact systemic treatment effects. Second, it is important to note that this study employed water bath heating to achieve mild hyperthermic tumor temperatures. While this approach does lead to uniform heating of the animal’s hind limb, heating is not constrained to the tumor [52]. Consequently, drug release occurs throughout the entire heated vasculature. Differences in heated tissue volume have been shown to affect the kinetics as well as total amount of intravascularly released drug, thus leading to varied amounts of free drug present in the systemic circulation [52]. The laser-based heating setup employed in this study provides more precise localized heating compared to water bath heating [36]. Nonetheless, combining thermosensitive liposomal vinorelbine with localized mild hyperthermia treatment via this heating setup resulted in reduced growth of the contralateral (i.e., unheated) tumor. This reduction in tumor growth could be attributed to systemically available thermosensitive liposomal vinorelbine and free vinorelbine. However, since these studies were performed in an immunocompetent animal model, it could also be due to immune mediated anti-tumor effects stimulated by mild hyperthermia and/or vinorelbine chemotherapy. While it is interesting to observe this anti-tumor effect at the contralateral site, it is important to note that this is a bilateral tumor model used to assess systemic anti-tumor capabilities. In general, metastatic tumor cell spread follows a highly complex process that is not reflected by bilateral tumor implantation [53]. Nonetheless, following primarily localized drug release of vinorelbine at the heated tumor we do observe a systemic anti-tumor effect. Thus, this study confirms the findings reported by Viglianti et al., albeit in a different animal model, and for a different thermosensitive liposome formulation.

### Immunocompetent animal models of rhabdomyosarcoma

We previously evaluated the combination of thermosensitive liposomal vinorelbine and mild hyperthermia in an immunocompromised murine model of RMS [39]. However, immunocompetent cancer models are necessary to evaluate the full potential of heat-triggered drug delivery approaches. While heating is used as the external trigger for drug release, it also elicits a series of effects that stimulate and enhance anti-tumor immune responses [26]. Most murine RMS models are based on human-derived xenografts and only a few syngeneic RMS animal models have been reported [54]. The M3–9-M RMS model employed was previously developed and characterized by Meadors et al. [35]. The authors demonstrated its immunogenicity and responsiveness to T-cell based immunotherapy [35]. Additionally, Highfill et al. demonstrated the role of PD-1 signaling in the immune escape of the M3–9-M tumors. Interestingly, in this study, PD-1 blockade therapy showed limited tumor growth inhibition effects when initiated later in the tumor development process [55]. Generally, orthotopic tumor models better recapitulate clinical disease, however, in the present study, tumors were grown subcutaneously due to limitations associated with the laser-based heating set-up [36,56–58].

### Previous studies combining thermosensitive liposomes and immune checkpoint inhibition

Kheirolomoom et al. have previously investigated similar strategies to increase the systemic treatment capabilities of thermosensitive liposomes. For example, the combination of a thermosensitive liposome formulation of doxorubicin with ultrasound mediated hyperthermia and intratumoral administration of CpG has previously been studied [59]. Here, CpG was employed as a local immune adjuvant to stimulate a more potent innate immune response. However, this therapy proved to be inefficient in treating distant (i.e., unheated) tumors. In a subsequent study, the authors showed that the addition of PD-1 blockade priming therapy to this treatment approach led to a potent anti-tumor T-cell response and successfully treated local as well as distant tumors [60]. The immunotherapy combination (i.e., i.t. CpG plus i.p. anti-PD-1) alone provided a potent systemic anti-tumor effect, without the addition of heat-triggered chemotherapy. Thus, based on this alone study, it is difficult to discern the contribution of thermosensitive liposomal doxorubicin to immunotherapy treatment. In our study, we also observed improvements in survival of animals receiving immunotherapy. Furthermore, we also see a significant survival benefit in animals treated with immunotherapy (i.e., anti-PD-1) in combination with thermosensitive liposomal vinorelbine and localized mild hyperthermia, compared to animals treated with immunotherapy alone. Albeit, in a different disease model and using a different treatment approach compared to the aforementioned study by Kheirolomoom et al. Ultimately, a comparison of the present study and the study conducted by Kheirolomoom et al highlights that the integration of immunotherapy into multimodal treatment regimens and evaluation of their efficacy in murine models of cancer is non-trivial.

### Mild hyperthermia and abscopal effects

As mentioned previously, the exact mechanism underlying the abscopal effect associated with localized therapies remains unclear [20]. However, it is generally recognized that a systemic cytotoxic T-cell mediated anti-tumor response plays a key role. Interestingly, mild hyperthermia has been shown to directly and indirectly influence components of the innate as well as adaptive immune response [61]. However, it is important to note that in most of these studies heat is applied for extended periods of time (≥ 1 h) and/or whole-body hyperthermia is employed instead of localized heating. In fact, very few *in vivo* studies have evaluated the potential abscopal effects that may be induced by treatment with localized mild hyperthermia alone. Mild hyperthermia is most commonly applied in combination with chemotherapy or radiotherapy treatment, thereby making it difficult to determine any effect of mild hyperthermia alone [62,63]. It was only recently that mild hyperthermia was combined with ICI to enhance abscopal effects [28,64]. Oei et al. evaluated hyperthermia alone and in combination with ICI in a highly metastatic mouse model of breast cancer (i.e., luciferase transfected 4T1 cells). The authors observed an increase in lung metastases following treatment with a combination of localized mild hyperthermia and ICI (i.e., anti-PD-1 and anti-CTLA-4). In contrast, Ibuki et al. employed the non-transfected 4T1 parent cell line in a bilateral subcutaneous tumor model. Here the combination of mild hyperthermia and ICI (i.e., anti-CTLA-4) led to strong tumor growth inhibition of both heated and unheated tumors [64]. The combination treatment also reduced metastatic spread to lungs (compared to ICI alone), resulting in an increase in overall survival. The contradictory results reported by Oei et al. and Ibuki et al. further highlight the need for more research into the combined effect of immunotherapy and mild hyperthermia [28,64]. While there are many differences in the study design and treatment strategies that might explain these results, there are four key differences that highlight the complexity of evaluating such a multimodal treatment approach. First, the immunogenic potential of the luciferase transfected cell line needs to be considered when evaluating an anti-tumor immune response [65]. Second, the anti-tumor immune response is heavily influenced and directed by the tumor microenvironment (i.e., orthotopic versus ectopic) [66]. Third, the immune effects mediated by mild hyperthermia treatment depend on the temperature and duration of heating as well as the specific heating technique [67,68]. And lastly, the timing of treatment relative to disease burden is critical when evaluating the efficacy of treatment induced anti-tumor immune effects. For example, the overall tumor burden has been shown to have an immunosuppressive effect [69]. In the present study, when PD-1 blockade monotherapy was initiated early (i.e., at a significantly smaller tumor volume) this led to complete tumor remission (albeit n = 2 mice), while delayed treatment resulted in limited tumor growth inhibition (**Figure S1**). This is in agreement with the previous work done by Highfill et al [55]. These studies demonstrate the complexity of this multimodal approach and highlight the need for detailed studies to better understand the parameters that influence treatment outcomes.

### Contributions of chemotherapy to systemic anti-tumor immune response

There is a growing body of evidence that suggests chemotherapy can exert a physiological response beyond a direct cytostatic or cytotoxic effect. In fact, some have proposed that the reason behind the success of certain chemotherapies lies in their ability to induce an anti-tumor immune response [70,71]. This is usually attributed to a specific form of regulated cell death, known as immunogenic cell death [72]. Indeed, several commonly used chemotherapeutic drugs have recently been identified as immunogenic cell death inducers. Among these are anthracyclines (e.g., doxorubicin) as well as microtubule inhibitors (e.g., taxanes and vinca alkaloids), including vinorelbine [73–75]. The classification of chemotherapeutic drugs as immunogenic cell death inducers is an active area of research. However, there is evidence to suggest that both localized mild hyperthermia as well as vinorelbine chemotherapy can create immunogenic tumors [76–82]. This would certainly be advantageous, and potentially synergistic, when administered in combination with ICI therapy. In the present study, such synergy is observed for the triple combination of thermosensitive liposomal vinorelbine, mild hyperthermia, and anti-PD-1. Specifically, this is demonstrated by enhanced efficacy at the distant tumor sites. Furthermore, this idea of exploiting the potentially immunostimulatory effects of vinorelbine is currently being investigated in clinical trials as a combination with immunotherapy (NCT03801304, NCT03518606, NCT04848454).

To conclude, thermosensitive liposomal vinorelbine plus localized mild hyperthermia in a bilateral tumor model of RMS was shown to have a significant anti-tumor effect at the heated primary site, resulting in tumor remission. Moreover, an effect at distant tumor sites was also observed for this treatment approach, albeit limited to reduced tumor growth rates. Interestingly, combining immune checkpoint inhibition therapy with thermosensitive liposomal vinorelbine and mild hyperthermia significantly improved treatment of distant tumor sites. Moreover, the addition of immune checkpoint inhibition therapy to localized mild hyperthermia alone produced a measurable abscopal effect. The underlying mechanism of this synergistic treatment effect will require further investigation. Yet, to the authors knowledge, this is the first study demonstrating that immune checkpoint inhibition therapy alone can enhance the systemic treatment capabilities of thermosensitive liposomes in combination with localized mild hyperthermia.

## Supporting information

Supplementary Informatonlementary

## 5. Acknowledgements

These studies were supported by a CIHR project grant to C.A. MR holds a Centre for Pharmaceutical Oncology scholarship. The authors acknowledge the use of equipment in the Centre for Pharmaceutical Oncology (CPO) at the University of Toronto as well as at STTARR Innovation Centre (University Health Network).

## 7. Abbreviations

CpG: CpG oligodeoxynucleotides
CTLA-4: cytotoxic T-lymphocyte-associated protein 4
DAMP: damage-associated molecular pattern
DPPC: 1,2-Dipalmitoyl-sn-glycero-3-phosphocholine
FBS: fetal bovine serum
HBS: HEPES buffered saline
HT: mild hyperthermia
i.p.: intraperitoneal
i.v.: intravenous
ICI: immune checkpoint inhibition
lyso-SPC: 1-stearoyl-2-lyso-sn-glycero-3-phosphocholine
Na_8_SOS: sodium sucrose octasulfate
NEAA: non-essential amino acid
P/S: penicillin and streptomycin
PD-1: programmed cell death protein 1
PEG_2k_-DSPE: N-(carbonyl-methoxypolyethyleneglycol 2000)-1,2-distearoyl-sn-glycero-3-phosphoethanolamine
HT: mild hyperthermia
RMS: rhabdomyosarcoma
TEA_8_SOS: sucrose octasulfate triethylammonium salt
ThermoVRL: thermosensitive liposomal vinorelbine
VRL: vinorelbine tartrate.

## Supplementary Information

**Figure S1:**
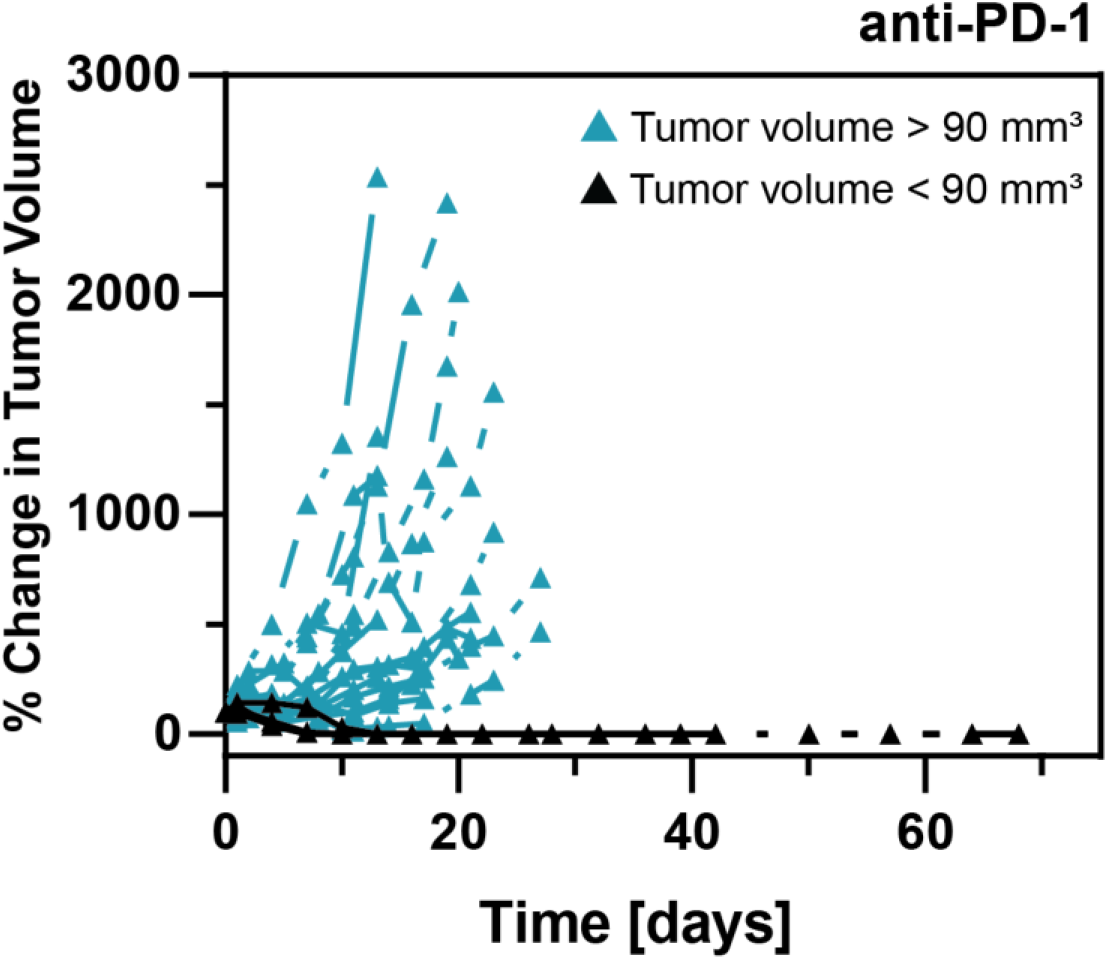
The % change in volume of subcutaneous M3–9-M tumors in male C57BL/6 mice that received PD-1 blockade therapy (i.e., anti-PD-1 antibody i.p. 24 hours prior to treatment day 0). PD-1 blockade therapy was administered twice per week for a total of 5 weeks. PD-1 blockade therapy led to complete remission when tumor volumes were < 90 mm³ on treatment day 0 (n = 2). In animals with tumor volumes > 90 mm³, PD-1 blockade therapy provided limited tumor growth inhibition (n = 8).

**Figure S2:**
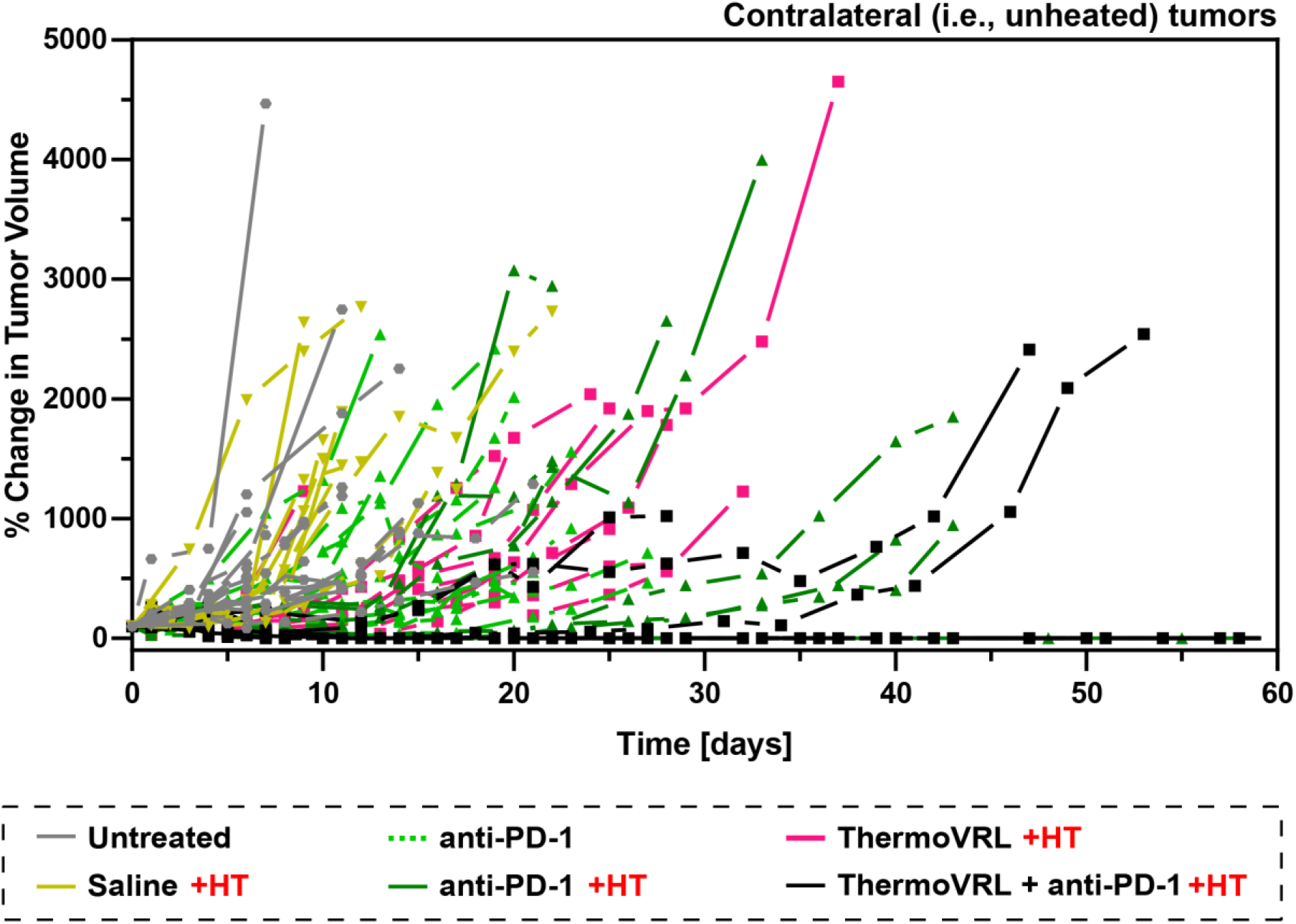
The % change in volume of contralateral (i.e., unheated) tumors in male C57BL/6 mice bearing bilateral, subcutaneous, M3–9-M tumors and receiving either no treatment (untreated), intraperitoneal PD-1 blockade therapy (anti-PD-1), intravenous saline (saline), or intravenous thermosensitive liposomal vinorelbine (ThermoVRL) at 10 mg VRL/kg body weight. As indicated, treatments were administered simultaneously with mild hyperthermia (+ HT; 42.5 °C, 25 min) localized to the primary tumor. n ≥ 8 per group.

**Figure S3:**
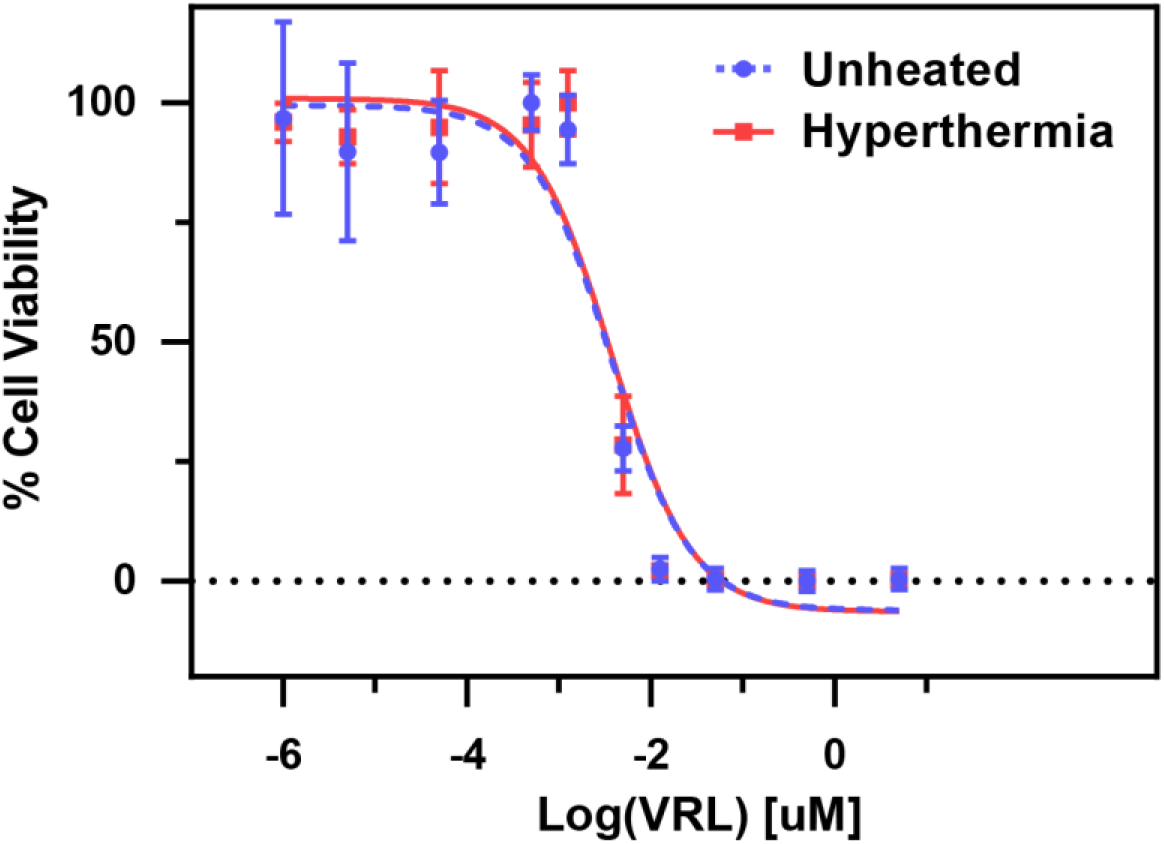
The in vitro cytotoxicity of VRL in M3–9-M cells with and without the addition of mild hyperthermia (42 °C for 1 h). Cells were treated with drug for a total incubation time of 72 h. The IC_50_ of VRL alone was found to be 3.8 ± 0.3 nM, and 3.7 ± 0.5 nM for VRL with the addition of mild hyperthermia (p > .05). Error bars represent SD of three independent experiments (n = 3).

## References

1. Crompton JG, Ogura K, Bernthal NM, Kawai A, Eilber FC. Local Control of Soft Tissue and Bone Sarcomas. J Clin Oncol. Wolters Kluwer; 2018;36:111–7.

2. Mahvi DA, Liu R, Grinstaff MW, Colson YL, Raut CP. Local Cancer Recurrence: The Realities, Challenges, and Opportunities for New Therapies. CA Cancer J Clin. 2018;68:488–505.

3. Mazzoleni S, Bisogno G, Garaventa A, Cecchetto G, Ferrari A, Sotti G, et al. Outcomes and prognostic factors after recurrence in children and adolescents with nonmetastatic rhabdomyosarcoma. Cancer. 2005;104:183–90.

4. Rhabdomyosarcoma - Childhood - Statistics [Internet]. Cancer.Net. 2012 [cited 2022 Jun 22]. Available from: https://www.cancer.net/cancer-types/rhabdomyosarcoma-childhood/statistics

5. M Fatih Okcu, John Hicks. Rhabdomyosarcoma in childhood, adolescence, and adulthood: Treatment [Internet]. UpToDate. 2022 [cited 2022 Jun 22]. Available from: https://www-uptodate-com.myaccess.library.utoronto.ca/contents/rhabdomyosarcoma-in-childhood-adolescence-and-adulthood-treatment?search=rhabdomyosarcoma%20treatment&source=search_result&selectedTitle=1~109&usage_type=default&display_rank=1#H31

6. Franco MS, Gomes ER, Roque MC, Oliveira MC. Triggered Drug Release From Liposomes: Exploiting the Outer and Inner Tumor Environment. Front Oncol [Internet]. 2021 [cited 2022 Mar 16];11. Available from: https://www.frontiersin.org/article/10.3389/fonc.2021.623760

7. Yatvin MB, Weinstein JN, Dennis WH, Blumenthal R. Design of Liposomes for Enhanced Local Release of Drugs by Hyperthermia. Science. American Association for the Advancement of Science; 1978;202:1290–3.

8. Motamarry A, Asemani D, Haemmerich D. Thermosensitive Liposomes. Liposomes [Internet]. 2017 [cited 2019 Nov 24]; Available from: https://www.intechopen.com/books/liposomes/thermosensitive-liposomes

9. Mannaris C, Efthymiou E, Meyre M-E, Averkiou MA. In Vitro Localized Release of Thermosensitive Liposomes with Ultrasound-Induced Hyperthermia. Ultrasound Med Biol. 2013;39:2011–20.

10. Viglianti BL, Dewhirst MW, Boruta RJ, Park J-Y, Landon C, Fontanella AN, et al. Systemic anti-tumour effects of local thermally sensitive liposome therapy. Int J Hyperth Off J Eur Soc Hyperthermic Oncol North Am Hyperth Group. 2014;30:385–92.

11. Regenold M, Bannigan P, Evans JC, Waspe A, Temple MJ, Allen C. Turning down the heat: The case for mild hyperthermia and thermosensitive liposomes. Nanomedicine Nanotechnol Biol Med. 2022;40:102484.

12. Lyon PC, Gray MD, Mannaris C, Folkes LK, Stratford M, Campo L, et al. Safety and feasibility of ultrasound-triggered targeted drug delivery of doxorubicin from thermosensitive liposomes in liver tumours (TARDOX): a single-centre, open-label, phase 1 trial. Lancet Oncol. 2018;19:1027–39.

13. de Maar JS, Suelmann BBM, Braat Mngja, van Diest PJ, Vaessen HHB, Witkamp AJ, et al. Phase I feasibility study of Magnetic Resonance guided High Intensity Focused Ultrasound-induced hyperthermia, Lyso-Thermosensitive Liposomal Doxorubicin and cyclophosphamide in de novo stage IV breast cancer patients: study protocol of the i-GO study. BMJ Open [Internet]. 2020 [cited 2021 Jan 8];10. Available from: https://www.ncbi.nlm.nih.gov/pmc/articles/PMC7692846/

14. Borys N, Dewhirst MW. Drug development of lyso-thermosensitive liposomal doxorubicin: Combining hyperthermia and thermosensitive drug delivery. Adv Drug Deliv Rev. 2021;178:113985.

15. Dillekås H, Rogers MS, Straume O. Are 90% of deaths from cancer caused by metastases? Cancer Med. 2019;8:5574–6.

16. Benzekry S, Tracz A, Mastri M, Corbelli R, Barbolosi D, Ebos JML. Modeling Spontaneous Metastasis following Surgery: An In Vivo-In Silico Approach. Cancer Res. 2016;76:535–47.

17. Mole RH. Whole Body Irradiation—Radiobiology or Medicine? Br J Radiol. The British Institute of Radiology; 1953;26:234–41.

18. Demaria S, Formenti SC. The abscopal effect 67 years later: from a side story to center stage. Br J Radiol. 2020;93:20200042.

19. Abuodeh Y, Venkat P, Kim S. Systematic review of case reports on the abscopal effect. Curr Probl Cancer. 2016;40:25–37.

20. Ngwa W, Irabor OC, Schoenfeld JD, Hesser J, Demaria S, Formenti SC. Using immunotherapy to boost the abscopal effect. Nat Rev Cancer. Nature Publishing Group; 2018;18:313–22.

21. Grass GD, Krishna N, Kim S. The immune mechanisms of abscopal effect in radiation therapy. Curr Probl Cancer. 2016;40:10–24.

22. Trommer M, Yeo SY, Persigehl T, Bunck A, Grüll H, Schlaak M, et al. Abscopal Effects in Radio-Immunotherapy—Response Analysis of Metastatic Cancer Patients With Progressive Disease Under Anti-PD-1 Immune Checkpoint Inhibition. Front Pharmacol [Internet]. 2019 [cited 2022 Jun 21];10. Available from: https://www.frontiersin.org/article/10.3389/fphar.2019.00511

23. Craig DJ, Nanavaty NS, Devanaboyina M, Stanbery L, Hamouda D, Edelman G, et al. The abscopal effect of radiation therapy. Future Oncol [Internet]. Future Medicine Ltd London, UK; 2021 [cited 2022 Jun 21]; Available from: https://www.futuremedicine.com/doi/10.2217/fon-2020-0994

24. Seiwert TY, Kiess AP. Time to Debunk an Urban Myth? The “Abscopal Effect” With Radiation and Anti– PD-1. J Clin Oncol. Wolters Kluwer; 2021;39:1–3.

25. Payne M, Bossmann SH, Basel MT. Direct treatment versus indirect: Thermo-ablative and mild hyperthermia effects. WIREs Nanomedicine Nanobiotechnology. 2020;12:e1638.

26. Skitzki JJ, Repasky EA, Evans SS. Hyperthermia as an immunotherapy strategy for cancer. Curr Opin Investig Drugs Lond Engl 2000. 2009;10:550–8.

27. Minnaar CA, Kotzen JA, Ayeni OA, Vangu M-D-T, Baeyens A. Potentiation of the Abscopal Effect by Modulated Electro-Hyperthermia in Locally Advanced Cervical Cancer Patients. Front Oncol. 2020;10:376.

28. Oei AL, Korangath P, Mulka K, Helenius M, Coulter JB, Stewart J, et al. Enhancing the abscopal effect of radiation and immune checkpoint inhibitor therapies with magnetic nanoparticle hyperthermia in a model of metastatic breast cancer. Int J Hyperth Off J Eur Soc Hyperthermic Oncol North Am Hyperth Group. 2019;36:47–63.

29. Zhou J, Wang G, Chen Y, Wang H, Hua Y, Cai Z. Immunogenic cell death in cancer therapy: Present and emerging inducers. J Cell Mol Med. 2019;23:4854–65.

30. Pfirschke C, Engblom C, Rickelt S, Cortez-Retamozo V, Garris C, Pucci F, et al. Immunogenic chemotherapy sensitizes tumors to checkpoint blockade therapy. Immunity. 2016;44:343–54.

31. Galluzzi L, Humeau J, Buqué A, Zitvogel L, Kroemer G. Immunostimulation with chemotherapy in the era of immune checkpoint inhibitors. Nat Rev Clin Oncol. Nature Publishing Group; 2020;17:725–41.

32. Marabelle A, Andtbacka R, Harrington K, Melero I, Leidner R, de Baere T, et al. Starting the fight in the tumor: expert recommendations for the development of human intratumoral immunotherapy (HIT-IT). Ann Oncol. 2018;29:2163–74.

33. Regenold M, Steigenberger J, Siniscalchi E, Dunne M, Casettari L, Heerklotz H, et al. Determining critical parameters that influence in vitro performance characteristics of a thermosensitive liposome formulation of vinorelbine. J Controlled Release [Internet]. 2020 [cited 2020 Sep 11]; Available from: http://www.sciencedirect.com/science/article/pii/S0168365920305009

34. Dou YN, Zheng J, Foltz WD, Weersink R, Chaudary N, Jaffray DA, et al. Heat-activated thermosensitive liposomal cisplatin (HTLC) results in effective growth delay of cervical carcinoma in mice. J Controlled Release. 2014;178:69–78.

35. Meadors JL, Cui Y, Chen Q-R, Song YK, Khan J, Merlino G, et al. Murine Rhabdomyosarcoma Is Immunogenic and Responsive to T-Cell-Based Immunotherapy. Pediatr Blood Cancer. 2011;57:921–9.

36. Dou YN, Weersink RA, Foltz WD, Zheng J, Chaudary N, Jaffray DA, et al. Custom-designed Laser-based Heating Apparatus for Triggered Release of Cisplatin from Thermosensitive Liposomes with Magnetic Resonance Image Guidance. J Vis Exp JoVE [Internet]. 2015 [cited 2020 Jan 18]; Available from: https://www.ncbi.nlm.nih.gov/pmc/articles/PMC4694025/

37. van Rhoon GC, Franckena M, ten Hagen TLM. A moderate thermal dose is sufficient for effective free and TSL based thermochemotherapy. Adv Drug Deliv Rev [Internet]. 2020 [cited 2020 Jul 30]; Available from: http://www.sciencedirect.com/science/article/pii/S0169409X2030020X

38. van Rhoon GC. Is CEM43 still a relevant thermal dose parameter for hyperthermia treatment monitoring? Int J Hyperthermia. Taylor & Francis; 2016;32:50–62.

39. Regenold M, Kan Kaneko, Xuehan Wang, H. Benson Peng, James C. Evans, Pauric Bannigan, et al. Triggered Release from Thermosensitive Liposomes Improves Tumor Targeting of Vinorelbine. Rev J Control Release. 2022;

40. Dunne M, Epp-Ducharme B, Sofias AM, Regenold M, Dubins DN, Allen C. Heat-activated drug delivery increases tumor accumulation of synergistic chemotherapies. J Controlled Release [Internet]. 2019 [cited 2019 Jul 8]; Available from: http://www.sciencedirect.com/science/article/pii/S0168365919303220

41. Kheirolomoom A, Lai C-Y, Tam SM, Mahakian LM, Ingham ES, Watson KD, et al. Complete regression of local cancer using temperature-sensitive liposomes combined with ultrasound-mediated hyperthermia. J Controlled Release. 2013;172:266–73.

42. Dou YN, Dunne M, Huang H, Mckee T, Chang MC, Jaffray DA, et al. Thermosensitive liposomal cisplatin in combination with local hyperthermia results in tumor growth delay and changes in tumor microenvironment in xenograft models of lung carcinoma. J Drug Target. 2016;24:865–77.

43. Needham D, Anyarambhatla G, Kong G, Dewhirst MW. A New Temperature-sensitive Liposome for Use with Mild Hyperthermia: Characterization and Testing in a Human Tumor Xenograft Model. Cancer Res. 2000;60:1197–201.

44. Bing C, Patel P, Staruch RM, Shaikh S, Nofiele J, Wodzak Staruch M, et al. Longer heating duration increases localized doxorubicin deposition and therapeutic index in Vx2 tumors using MR-HIFU mild hyperthermia and thermosensitive liposomal doxorubicin. Int J Hyperthermia. Taylor & Francis; 2019;36:195–202.

45. Gasselhuber A, Dreher MR, Negussie A, Wood BJ, Rattay F, Haemmerich D. Mathematical spatiotemporal model of drug delivery from low temperature sensitive liposomes during radiofrequency tumour ablation. Int J Hyperthermia. Taylor & Francis; 2010;26:499–513.

46. Ranjan A, Jacobs G, Woods DL, Negussie AH, Partanen A, Yarmolenko PS, et al. Image-guided drug delivery with magnetic resonance guided high intensity focused ultrasound and temperature sensitive liposomes in a rabbit Vx2 tumor model. J Controlled Release. 2012;158:487–94.

47. Kok HP, Cressman ENK, Ceelen W, Brace CL, Ivkov R, Grüll H, et al. Heating technology for malignant tumors: a review. Int J Hyperthermia. 2020;37:711–41.

48. Santos MA, Wu S-K, Regenold M, Allen C, Goertz DE, Hynynen K. Novel fractionated ultrashort thermal exposures with MRI-guided focused ultrasound for treating tumors with thermosensitive drugs. Sci Adv. American Association for the Advancement of Science; 2020;6:eaba5684.

49. Crezee J, Zweije R, Sijbrands J, Kok HP. Dedicated 70 MHz RF systems for hyperthermia of challenging tumor locations. Int J Microw Wirel Technol. Cambridge University Press; 2020;12:839–47.

50. World Health Organization. Cancer-WHO [Internet]. World Health Organ. 2022 [cited 2022 May 16]. Available from: https://www.who.int/news-room/fact-sheets/detail/cancer

51. Zhou J, Lu X, Chang W, Wan C, Lu X, Zhang C, et al. PLUS: Predicting cancer metastasis potential based on positive and unlabeled learning. PLOS Comput Biol. Public Library of Science; 2022;18:e1009956.

52. Untargeted Large Volume Hyperthermia Reduces Tumor Drug Uptake From Thermosensitive Liposomes. IEEE Open J Eng Med Biol. 2021;2:187–97.

53. Fares J, Fares MY, Khachfe HH, Salhab HA, Fares Y. Molecular principles of metastasis: a hallmark of cancer revisited. Signal Transduct Target Ther. Nature Publishing Group; 2020;5:1–17.

54. Nakahata K, Simons BW, Pozzo E, Shuck R, Kurenbekova L, Prudowsky Z, et al. K-Ras and p53 mouse model with molecular characteristics of human rhabdomyosarcoma and translational applications. Dis Model Mech. 2022;15:dmm049004.

55. Highfill SL, Cui Y, Giles AJ, Smith JP, Zhang H, Morse E, et al. Disruption of CXCR2-Mediated MDSC Tumor Trafficking Enhances Anti-PD1 Efficacy. Sci Transl Med. American Association for the Advancement of Science; 2014;6:237ra67–237ra67.

56. Guerin MV, Finisguerra V, Van den Eynde BJ, Bercovici N, Trautmann A. Preclinical murine tumor models: A structural and functional perspective. Settleman J, Kawakami Y, editors. eLife. eLife Sciences Publications, Ltd; 2020;9:e50740.

57. Zhang W, Fan W, Rachagani S, Zhou Z, Lele SM, Batra SK, et al. Comparative Study of Subcutaneous and Orthotopic Mouse Models of Prostate Cancer: Vascular Perfusion, Vasculature Density, Hypoxic Burden and BB2r-Targeting Efficacy. Sci Rep. Nature Publishing Group; 2019;9:11117.

58. Brand M, Laban S, Theodoraki M-N, Doescher J, Hoffmann TK, Schuler PJ, et al. Characterization and Differentiation of the Tumor Microenvironment (TME) of Orthotopic and Subcutaneously Grown Head and Neck Squamous Cell Carcinoma (HNSCC) in Immunocompetent Mice. Int J Mol Sci. 2020;22:247.

59. Kheirolomoom A, Ingham ES, Mahakian LM, Tam SM, Silvestrini MT, Tumbale SK, et al. CpG expedites regression of local and systemic tumors when combined with activatable nanodelivery. J Control Release Off J Control Release Soc. 2015;220:253–64.

60. Kheirolomoom A, Silvestrini MT, Ingham ES, Mahakian LM, Tam SM, Tumbale SK, et al. Combining activatable nanodelivery with immunotherapy in a murine breast cancer model. J Controlled Release. 2019;303:42–54.

61. Frey B, Weiss E-M, Rubner Y, Wunderlich R, Ott OJ, Sauer R, et al. Old and new facts about hyperthermia-induced modulations of the immune system. Int J Hyperthermia. 2012;28:528–42.

62. Dunne M, Regenold M, Allen C. Hyperthermia can alter tumor physiology and improve chemo- and radio-therapy efficacy. Adv Drug Deliv Rev [Internet]. 2020 [cited 2020 Jul 27]; Available from: http://www.sciencedirect.com/science/article/pii/S0169409X20300831

63. Issels RD, Lindner LH, von Bergwelt-Baildon M, Lang P, Rischpler C, Diem H, et al. Systemic antitumor effect by regional hyperthermia combined with low-dose chemotherapy and immunologic correlates in an adolescent patient with rhabdomyosarcoma – a case report. Int J Hyperthermia. Taylor & Francis; 2020;37:55–65.

64. Ibuki Y, Takahashi Y, Tamari K, Minami K, Seo Y, Isohashi F, et al. Local hyperthermia combined with CTLA-4 blockade induces both local and abscopal effects in a murine breast cancer model. Int J Hyperthermia. Taylor & Francis; 2021;38:363–71.

65. Baklaushev VP, Kilpeläinen A, Petkov S, Abakumov MA, Grinenko NF, Yusubalieva GM, et al. Luciferase Expression Allows Bioluminescence Imaging But Imposes Limitations on the Orthotopic Mouse (4T1) Model of Breast Cancer. Sci Rep. 2017;7:7715.

66. Mariana Varna, Philippe Bertheau, Luc G. Legrès. Tumor Microenvironment in Human Tumor Xenografted Mouse Models. J Anal Oncol [Internet]. 2014 [cited 2022 Jul 5];3. Available from: https://neoplasiaresearch.com/pms/index.php/jao/article/view/226

67. Beachy SH, Repasky EA. Toward establishment of temperature thresholds for immunological impact of heat exposure in humans. Int J Hyperth Off J Eur Soc Hyperthermic Oncol North Am Hyperth Group. 2011;27:344–52.

68. Baronzio GF, Seta RD, D’Amico M, Baronzio A, Freitas I, Forzenigo G, et al. Effects of Local and Whole Body Hyperthermia on Immunity [Internet]. Madame Curie Biosci. Database Internet. Landes Bioscience; 2013 [cited 2022 Jun 13]. Available from: http://www.ncbi.nlm.nih.gov/books/NBK6083/

69. Kim SI, Cassella CR, Byrne KT. Tumor Burden and Immunotherapy: Impact on Immune Infiltration and Therapeutic Outcomes. Front Immunol [Internet]. 2021 [cited 2022 Jul 5];11. Available from: https://www.frontiersin.org/articles/10.3389/fimmu.2020.629722

70. Hiam-Galvez KJ, Allen BM, Spitzer MH. Systemic immunity in cancer. Nat Rev Cancer. Nature Publishing Group; 2021;21:345–59.

71. Galluzzi L, Buqué A, Kepp O, Zitvogel L, Kroemer G. Immunological Effects of Conventional Chemotherapy and Targeted Anticancer Agents. Cancer Cell. 2015;28:690–714.

72. Lin RA, Lin JK, Lin S-Y. Mechanisms of immunogenic cell death and immune checkpoint blockade therapy. Kaohsiung J Med Sci. 2021;37:448–58.

73. Bezu L, Sauvat A, Humeau J, Gomes-da-Silva LC, Iribarren K, Forveille S, et al. eIF2α phosphorylation is pathognomonic for immunogenic cell death. Cell Death Differ. Nature Publishing Group; 2018;25:1375–93.

74. Roselli M, Cereda V, di Bari MG, Formica V, Spila A, Jochems C, et al. Effects of conventional therapeutic interventions on the number and function of regulatory T cells. OncoImmunology. Taylor & Francis; 2013;2:e27025.

75. Wen C-C, Chen H-M, Chen S-S, Huang L-T, Chang W-T, Wei W-C, et al. Specific microtubule-depolymerizing agents augment efficacy of dendritic cell-based cancer vaccines. J Biomed Sci. 2011;18:44.

76. Orecchioni S, Talarico G, Labanca V, Calleri A, Mancuso P, Bertolini F. Vinorelbine, cyclophosphamide and 5-FU effects on the circulating and intratumoural landscape of immune cells improve anti-PD-L1 efficacy in preclinical models of breast cancer and lymphoma. Br J Cancer. 2018;118:1329–36.

77. Yu Q, Tang X, Zhao W, Qiu Y, He J, Wan D, et al. Mild hyperthermia promotes immune checkpoint blockade-based immunotherapy against metastatic pancreatic cancer using size-adjustable nanoparticles. Acta Biomater. 2021;133:244–56.

78. Hurwitz MD. Hyperthermia and immunotherapy: clinical opportunities. Int J Hyperthermia. Taylor & Francis; 2019;36:4–9.

79. Altinoz MA, Ozpinar A, Alturfan EE, Elmaci I. Vinorelbine’s anti-tumor actions may depend on the mitotic apoptosis, autophagy and inflammation: hypotheses with implications for chemo-immunotherapy of advanced cancers and pediatric gliomas. J Chemother. Taylor & Francis; 2018;30:203–12.

80. Moskowitz AJ, Shah G, Schöder H, Ganesan N, Drill E, Hancock H, et al. Phase II Trial of Pembrolizumab Plus Gemcitabine, Vinorelbine, and Liposomal Doxorubicin as Second-Line Therapy for Relapsed or Refractory Classical Hodgkin Lymphoma. J Clin Oncol. Wolters Kluwer; 2021;39:3109–17.

81. D’Ascanio M, Pezzuto A, Fiorentino C, Sposato B, Bruno P, Grieco A, et al. Metronomic Chemotherapy with Vinorelbine Produces Clinical Benefit and Low Toxicity in Frail Elderly Patients Affected by Advanced Non-Small Cell Lung Cancer. BioMed Res Int. Hindawi; 2018;2018:e6278403.

82. Vergnenegre A, Monnet I, Bizieux A, Bernardi M, Chiapa AM, Léna H, et al. Open-label Phase II trial to evaluate safety and efficacy of second-line metronomic oral vinorelbine–atezolizumab combination for stage-IV non-small-cell lung cancer – VinMetAtezo trial, (GFPC‡ 04-2017). Future Oncol. Future Medicine; 2020;16:5–10.

