## Supplementary Informatonlementary for "Harnessing Immunotherapy to Enhance the Systemic Anti-Tumor Effects of Thermosensitive Liposomes"

### Supplementary Information

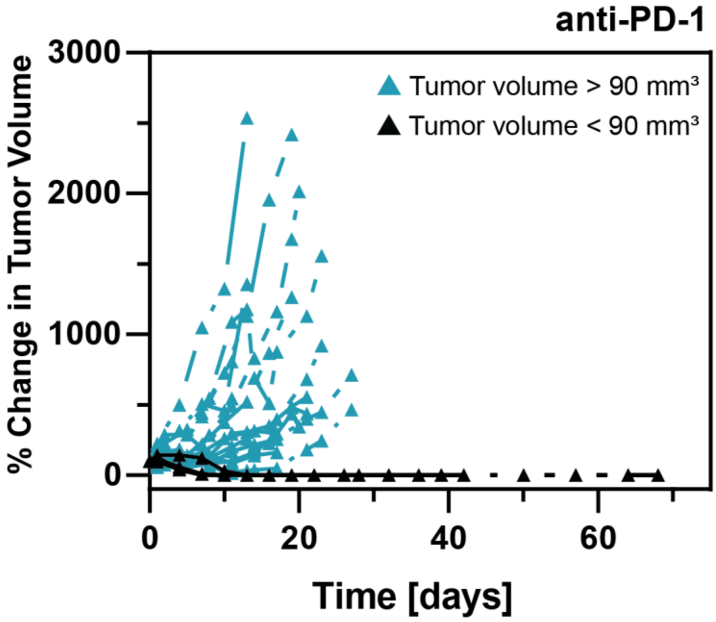


**Figure S1:** The % change in volume of subcutaneous M3–9-M tumors in male C57BL/6 mice that received PD-1 blockade therapy (i.e., anti-PD-1 antibody i.p. 24 hours prior to treatment day 0). PD-1 blockade therapy was administered twice per week for a total of 5 weeks. PD-1 blockade therapy led to complete remission when tumor volumes were < 90 $mm^3$ on treatment day 0 (n = 2). In animals with tumor volumes > 90 $mm^3$, PD-1 blockade therapy provided limited tumor growth inhibition (n = 8).

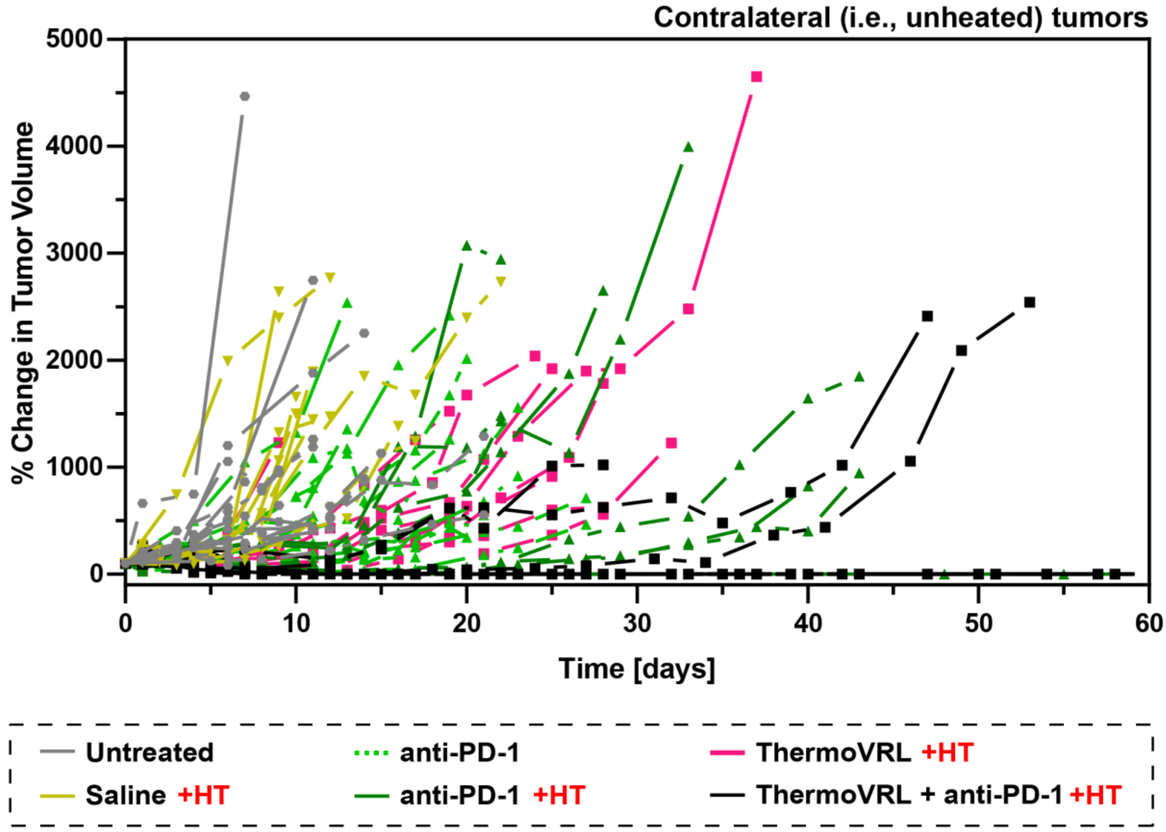


**Figure S2:** The % change in volume of contralateral (i.e., unheated) tumors in male C57BL/6 mice bearing bilateral, subcutaneous, M3–9-M tumors and receiving either no treatment (untreated), intraperitoneal PD-1 blockade therapy (anti-PD-1), intravenous saline (saline), or intravenous thermosensitive liposomal vinorelbine (ThermoVRL) at 10 mg VRL/kg body weight. As indicated, treatments were administered simultaneously with mild hyperthermia (+ HT; 42.5 °C, 25 min) localized to the primary tumor. n ≥ 8 per group.

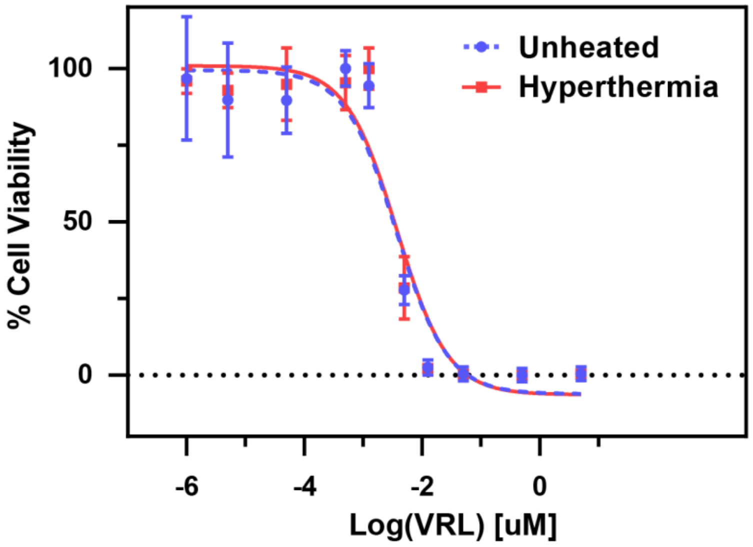


**Figure S3:** The in vitro cytotoxicity of VRL in M3–9-M cells with and without the addition of mild hyperthermia (42 °C for 1 h). Cells were treated with drug for a total incubation time of 72 h. The $IC_{50}$ of VRL alone was found to be 3.8 ± 0.3 nM, and 3.7 ± 0.5 nM for VRL with the addition of mild hyperthermia ($p > .05$). Error bars represent SD of three independent experiments (n = 3).
